# Single-cell multi-omics analyses reveal EZH2 as a main driver of retinoic acid resistance in PLZF-RARA leukemia

**DOI:** 10.1101/2022.01.04.474890

**Authors:** M. Poplineau, N. Platet, A. Mazuel, L. Hérault, S. Koide, W. Kuribayashi, N. Carbuccia, L. N’Guyen, J. Vernerey, M. Oshima, D. Birnbaum, A. Iwama, E. Duprez

## Abstract

Cancer relapse is caused by a subset of malignant cells that are resistant to treatment. To characterize resistant cells and their vulnerabilities, we studied the retinoic acid (RA)-resistant PLZF-RARA acute promyelocytic leukemia (APL) using single-cell multi-omics. We uncovered transcriptional and chromatin heterogeneity in leukemia cells and identified a subset of cells resistant to RA that depend on a fine-tuned transcriptional network targeting the epigenetic regulator Enhancer of Zeste Homolog 2 (EZH2). Epigenomic and functional analyses validated EZH2 selective dependency of PLZF-RARA leukemia and its driver role in RA resistance. Targeting pan-EZH2 activities (canonical/non-canonical) was necessary to eliminate leukemia relapse initiating cells, which underlies a dependency of resistant cells on an EZH2 non-canonical activity and the necessity to degrade EZH2 to overcome resistance.

Our study provides critical insights into the mechanisms of RA resistance that allow us to eliminate treatment-resistant leukemia cells by targeting EZH2, thus highlighting a potential targeted therapy approach.

**HIGHLIGHTS:**

- sc-RNAseq identifies PLZF-RARA leukemia heterogeneity and retinoic acid resistant cells
- sc-ATACseq refines leukemic cell identity and resolves retinoic acid resistant networks
- EZH2 is a selective dependency of PLZF-RARA leukemia and drives retinoic acid resistance
- Targeting pan-EZH2 activities (canonical/non-canonical) is necessary to overcome leukemia onset

## INTRODUCTION

Acute myeloid leukemia (AML), characterized by clonal growth and evolution of an undifferentiated hematopoietic cell is a heterogeneous disease in terms of cell-of-origin, genetic and epigenetic alterations, clinical features and treatment outcomes^1^. Yet, recurrent chromosomal rearrangements leading to the generation of oncogenic fusion proteins are common in AML and an abundant literature has emphasized their driver role in its development and response to therapeutics ^2^.

Acute promyelocytic leukemia (APL) is a class of AML that accounts for 10–15% of all cases and is characterized by recurrent chromosomal translocations involving invariably the gene encoding the Retinoic Acid Receptor alpha (RARA) (17q21) with several fusion partners, such as *PML* (15q22) or *PLZF* (11q23) (For review ^3^). The resulting X-RARA fusion proteins were among the first transcription factors (TFs) to be identified as drivers of cancer ^4^. They behave as RARA signaling repressors due to their ability to oligomerize and to recruit epigenetic repressors at cis-regulatory DNA regions of RARA target genes and initiate oncogenic gene expression signatures ^5^.

APL patients with PML-RARA fusion, are exquisitely sensitive to Retinoic Acid (RA) treatment, a sensitivity that has made APL one of the most successful examples of targeted therapy to mutational events^6; 7^. Although very successfully advanced, RA-targeted-treatment of APL still poses clinical challenges. Numerous mechanisms have been proposed to explain inherent and acquired RA resistance in leukemia including epigenetic mechanisms linked to RA signaling ^8; 9^.

APL patients with t(11;17) bearing the PLZF-RARA fusion protein respond poorly to RA and remain clinically resistant ^10^. To identify actionable vulnerabilities that will prevent resistance and facilitate treatment response, much work has been done to understand the cellular and molecular bases of PLZF-RARA APL physiopathology ^11^. Contrary to PML-RARA APL, pharmacological doses of RA, although inducing partial differentiation of the PLZF-RARA blasts, do not clear the Leukemia Initiating cell (LIC) of this APL variant ^12; 13^. This highlighted the uncoupling between blast differentiation and tumor eradication. The PLZF moiety of the fusion is thought to play a determinant role in RA-resistance. PLZF is a potent transcriptional repressor that can interact by itself with epigenetic complexes ^14; 15^. Given that PLZF binding sites with corepressors are conserved in the PLZF-RARA fusion, one hypothesis is that inappropriate recruitment of functional epigenetic repressors, such as the Polycomb Repressor Complex 1 (PRC1), would trigger epigenetic imbalance at very specific genomic loci in an RA insensitive manner ^16^. This is consistent with the observed degradation of PLZF-RARA under RA treatment ^17^ suggesting a persistent mechanism even in the absence of detectable fusion involving chromatin modifications. However, the molecular basis for the resistance of PLZF-RARA expressing cells and why some cells retain their self-renewal capacity while others do not after RA treatment remain unknown.

To advance in the understanding of RA resistance, we studied APL cellular heterogeneity by integrating scRNA-seq and scATAC-seq data obtained from PLZF-RARA transformed promyelocytes treated with RA. Establishing cellular clusters and arranging them in hierarchies helped us to identify a subpopulation of transformed promyelocytes that were insensitive to RA-induced differentiation and characterized by a DNA repair and replication gene expression signature and high expression of Enhancer of Zeste Homolog 2 (EZH2), the catalytic subunit of Polycomb Repressive Complex 2 (PRC2). Because EZH2 contribution to the pathogenesis of leukemia is contrasted ^18^, we further explored EZH2 function in APL development and RA treatment response. We discovered a dual role of EZH2 in APL onset and RA response, suggesting the need to target the non-histone methyltransferase activity of EZH2 for leukemia clearance.

## RESULTS

### Single-cell transcriptome analysis identifies a subset of retinoic acid resistant leukemic cells with DNA repair, replication and proliferation signatures

To decipher mechanisms linked to APL t(11;17) relapse and identify features of RA-resistant leukemic cells, we used droplet-based single-cell RNA sequencing (scRNA-seq) on the 10X Genomics platform for the PLZF-RARA leukemic cells obtained from bone marrow of PLZF-RARA-TG transplanted mice at day 17 post transplantation, treated or not with RA (**Figure S1A**). Effect of the RA-treatment was monitored by following increased expression of cell surface markers (Cd11b and Gr1) and by morphology analyses of FACS-sorted promyelocytes (**Figures S1B-S1C**). We obtained the transcriptome of 5106 untreated (NT) and 6794 RA-treated (d7) promyelocytes. Samples were then integrated to reduce batch effect and cell cycle genes were regressed using the Seurat workflow ^19^. Dimensionality reduction and unsupervised clustering in all cells (treated and untreated), identified six clusters that were annotated by enrichment analysis of their gene markers and visualized with uniform manifold approximation and projection (UMAP) ^20^ (**Figures 1A-B, S1D**). Five of the clusters had neutrophil associated signatures (Prom1, Prom2, Prom3, NeuRA1, NeuRA2) consistent with the promyelocytic stage of induced leukemia, while one cluster was characterized by genes involved in DNA replication associated with DNA repair and proliferation processes (ReP) (**Figure 1B, S1E, Tables S1A-C**). Prom1, Prom2 and Prom3 were enriched with untreated cells suggesting their fading upon RA treatment (**Figure 1C**), while NeuRA1 and NeuRA2 appeared almost exclusively in RA-treated cells. The terminal myeloid differentiation status of these two groups was supported by their gene signature (**Figure S1F**). By contrast the ReP cluster was characterized by the expression of genes involved in homologous recombination and DNA replication such as *Rad51, Pcna* and *Mcm3* (**Figure S1G**) and was equally composed of NT and RA-treated cells (**Figure 1C**) suggesting that RA treatment did not impact on the cell identity of these promyelocytes.

**Figure 1:**
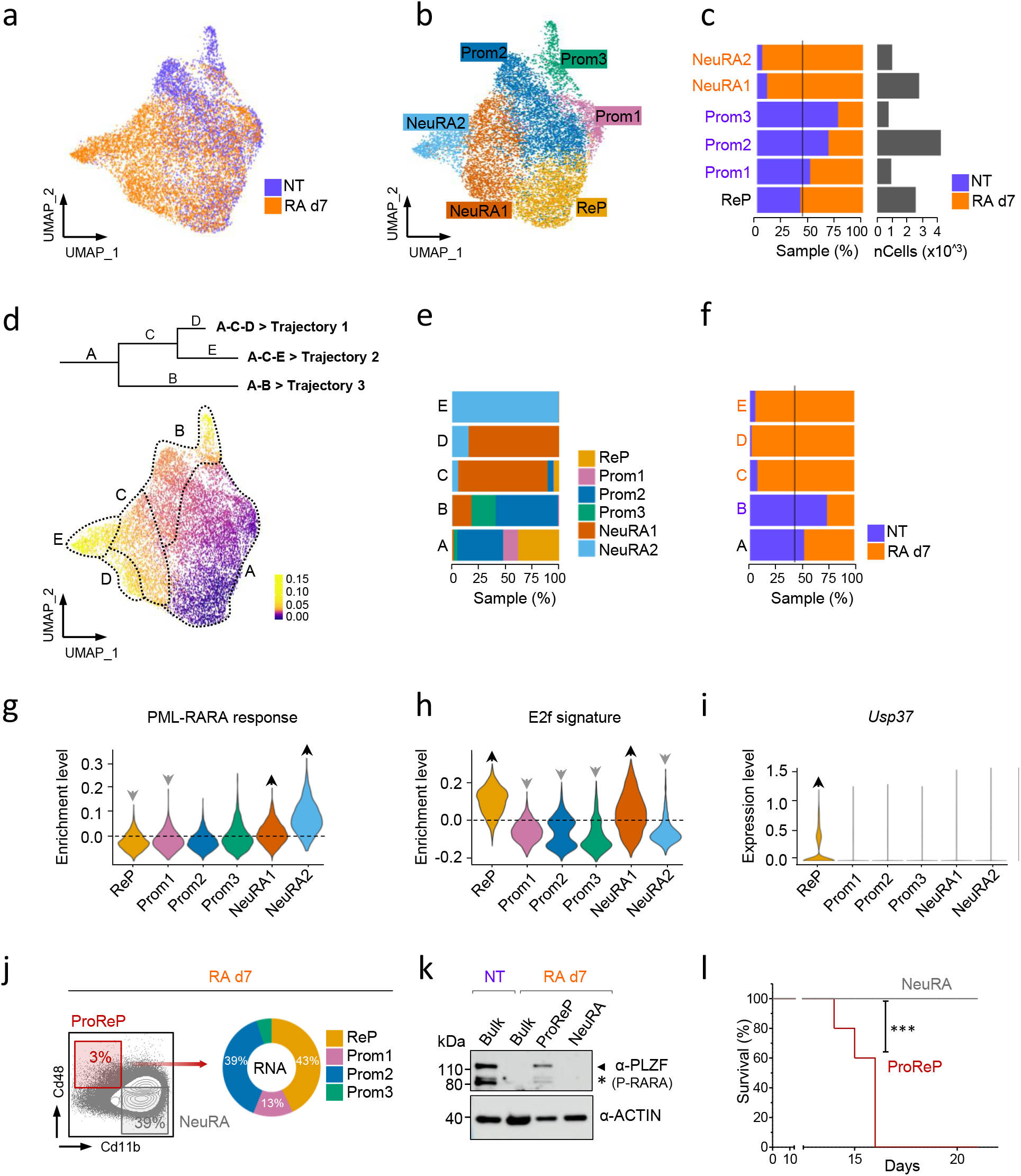
Single-cell transcriptome analysis identifies a subset of retinoic acid resistant PLZF-RARA leukemic cells with a specific signature. (**A**) UMAP visualization of the scRNA-seq dataset colored by condition: NT and RA d7 (total integrated cells: 11,900). (**B**) UMAP visualization of NT and RA d7 cells colored by cluster. Six clusters were identified by unsupervised clustering and characterized with differential gene expression and gene set enrichment analysis. ReP, DNA repair/Replication/Proliferation; Prom1-3, Neutrophil 1-3; NeuRA1-2, Neutrophil Retinoic Acid 1-2. (**C**) Left panel: NT and RA cell distribution in each cluster. The black vertical line indicates expected NT and RA cell proportions according to the dataset size. Names of the clusters for which the proportion of NT or RA cells was significantly higher than expected (hypergeometric test p-value <0.01) are colored in purple for NT cells and in orange for RA cells. Right panel: absolute cell number per cluster. (**D**) Differentiation trajectory obtained using STREAM. On top, schematic representation of the different states (A, B, C, D, E) in each trajectory (Trajectory 1: A-C-D, Trajectory 2: A-C-E, Trajectory 3: A-B). Below, UMAP is colored according to the pseudotime value of each cell. Color scale was rescaled to better fit the score distribution. (**E**) Barplot showing the cluster distribution in each state of the pseudotime. (**F**) NT and RA cell distribution in each state of the pseudotime. Names of the clusters for which the proportion of NT or RA cells was significantly higher than expected (hypergeometric test *p* value <0.01) are colored in purple for NT cells and in orange for RA cells. (**G**) Violin plot showing PML-RARA response signature scores per cluster. Signature score represents the global expression of annotated genes for the selected signature. (**H**) Violin plot showing E2f signature in each cluster. The signature score represents the global expression of annotated genes for the E2f signature. For **G** and **H** The gray/black arrows pointing down/up indicate significant lower/higher expression in the cluster against all the other clusters (average score difference > 0.02 and p-value adjusted < 0.05). (**I**) Violin plot showing *Usp37* expression in each cluster. Black arrows pointing up indicates significant higher expression in the cluster against all the other clusters (average log2|FC|> 0.1 and p-value adjusted < 0.05). (**J**) Gating strategy for isolating ProReP cells (Prom1-3 and ReP, C-Kit^+^, Gr1^+^, Cd48^+^, Cd11b^-^) and NeuRA (NeuRA1-2 : C-Kit^+^, Gr1^+^, Cd48^-^, Cd11b^+^). The percentage represents the proportion of ProReP and NeuRA in the promyelocyte population according to the FACS analysis. The donut plot represents the proportion of Prom1-3 and ReP cells into the ProReP population according to the scRNA-seq analysis. (**K**) PLZF-RARA (black arrow: full length, black star: degraded form) detected by western blotting in untreated (NT) and treated promyelocytes (RA d7) indicate population. Actin is used as loading control. (**L**) Survival rate (Kaplan Meier survival analysis, https://www.statskingdom.com/kaplan-meier.html) of mice transplanted with ProReP or NeuRA cells. *** p-value < 0.001.

To determine the potential differentiation journey of the RA-treated transformed promyelocytes, we ordered on pseudotime NT and RA-treated cells based on their transcriptional similarities with the Single-cell Trajectories Reconstruction, Exploration And Mapping (STREAM) pipeline ^21^ (**Figures 1D-F**). We revealed three trajectories split into five different states of promyelocytes (segments A, B, C, D, E) (**Figure 1D**) and defined the departure of the trajectories at the extremity of state A, which was mostly composed of ReP cells. Trajectories 1 (states A-C-D) and 2 (states A-C-E) ended towards RA differentiated cells, since their final states C, D and E were largely composed of RA-treated cells grouped in NeuRA1 and NeuRA2, respectively (**Figures 1E-F**). This data confirmed a pronounced differentiating effect of RA on a portion of PLZF-RARA expressing cells consistently with a stronger expression of differentiation marker in NeuRA2 (**Figure S1F**). Interestingly, the third trajectory (state B), which went far into the pseudotime was composed mostly of NT cells grouped in Prom2 and Prom3 (**Figures 1E-F**), suggesting a spontaneous differentiation program in leukemic cells. This analysis showed a pronounced but partial differentiating effect of RA on PLZF-RARA expressing cells and designated cells in the ReP cluster as the treatment-persistent cells.

To further investigate the RA (un)responsiveness of the identified clusters, we took advantage of available transcriptional signatures reflecting RA sensitivity of PML-RARA *versus* RA resistant PLZF-RARA murine APL ^13^. We found that the NeuRA1 and NeuRA2 clusters highly expressed a computed PML-RARA RA-responsive signature (see Mat & Med) confirming that the two clusters were composed of RA-responsive blasts (**Figure 1G**). By contrast, the ReP cluster was highly enriched with the proliferative E2F signature (**Figure 1H**), which persisted upon RA treatment (**Figure S1H, upper panel**), likely supporting the malignancy and RA insensitivity of these cells. Interestingly, the ReP cluster specifically expressed the deubiquitinase *Usp37* involved in PLZF-RARA fusion stability ^22^ and this expression was not impacted by the RA treatment (**Figures 1I, S1H, lower panel**). We attempted to purify the ReP cluster after RA treatment based on the expression of specific surface markers identified in our scRNA-seq dataset. Taking into account the expression of the resolving cell surface markers Cd48 and Itgam (Cd11b), we were limited to isolate Prom1-3 and ReP (ProReP) from NeuRA1 and NeuRA2 (NeuRA) cells (**Figures 1J, S1I**). After RA treatment, whereas the PLZF-RARA fusion was neither detected in the bulk nor in NeuRA populations, ProReP cells did maintain PLZF-RARA expression (**Figure 1K**) and kept the potential to develop leukemia in transplanted mice (**Figure 1L**).

Altogether, these results reveal transcriptional heterogeneity and differentiation states within the PLZF-RARA transformed cells and identify the ReP cluster, which exhibits no differentiation features, high E2F signature and PLZF-RARA residual expression, as the potential driver of RA resistance of PLZF-RARA leukemia.

### Single-cell integrative multi-omics analysis highlights chromatin genes as responsible for RA resistance in PLZF-RARA expressing cells

To characterize at the regulatory/epigenomic level the heterogeneity underlying PLZF-RARA transformation and RA response, we generated a chromatin accessibility profile (sc-ATACseq) of 4,215 RA-treated cells and 5,806 untreated cells in our PLZF-RARA TG mouse model. To link transcriptome variations with changes in epigenome and to better define cell identity and function, we first imputed gene activity from the ATAC signal in order to map cells from our scATAC-seq data to the six scRNA-seq defined clusters using pairwise correspondences ^23^. We thus defined a new shared low-dimensional space (**Figures 2A-B, S2A**) in which we showed an overall good concordance between the two levels of information (scATAC-seq and scRNA-seq), emphasizing the link between chromatin structure and transcription (**Figures 2A-B)**. This concordance was especially high on RA-treated promyelocytes (**Figures 2A-B)**, confirming the known effect of RA on differentiation through remodeling of chromatin landscape ^24^. Consistent with our scRNA-seq data, the ratio of scATAC-seq cells in the ReP cluster was not affected by RA treatment (7% in both conditions). The low impact of RA on chromatin opening in the ReP cluster in comparison to other clusters was confirmed by Coverage Plot visualization at some selected genes (**Figure 2C**). Thus, this analysis supports the important role of chromatin in RA response and confirms the little impact of RA on the ReP cluster at both RNA and chromatin levels.

**Figure 2:**
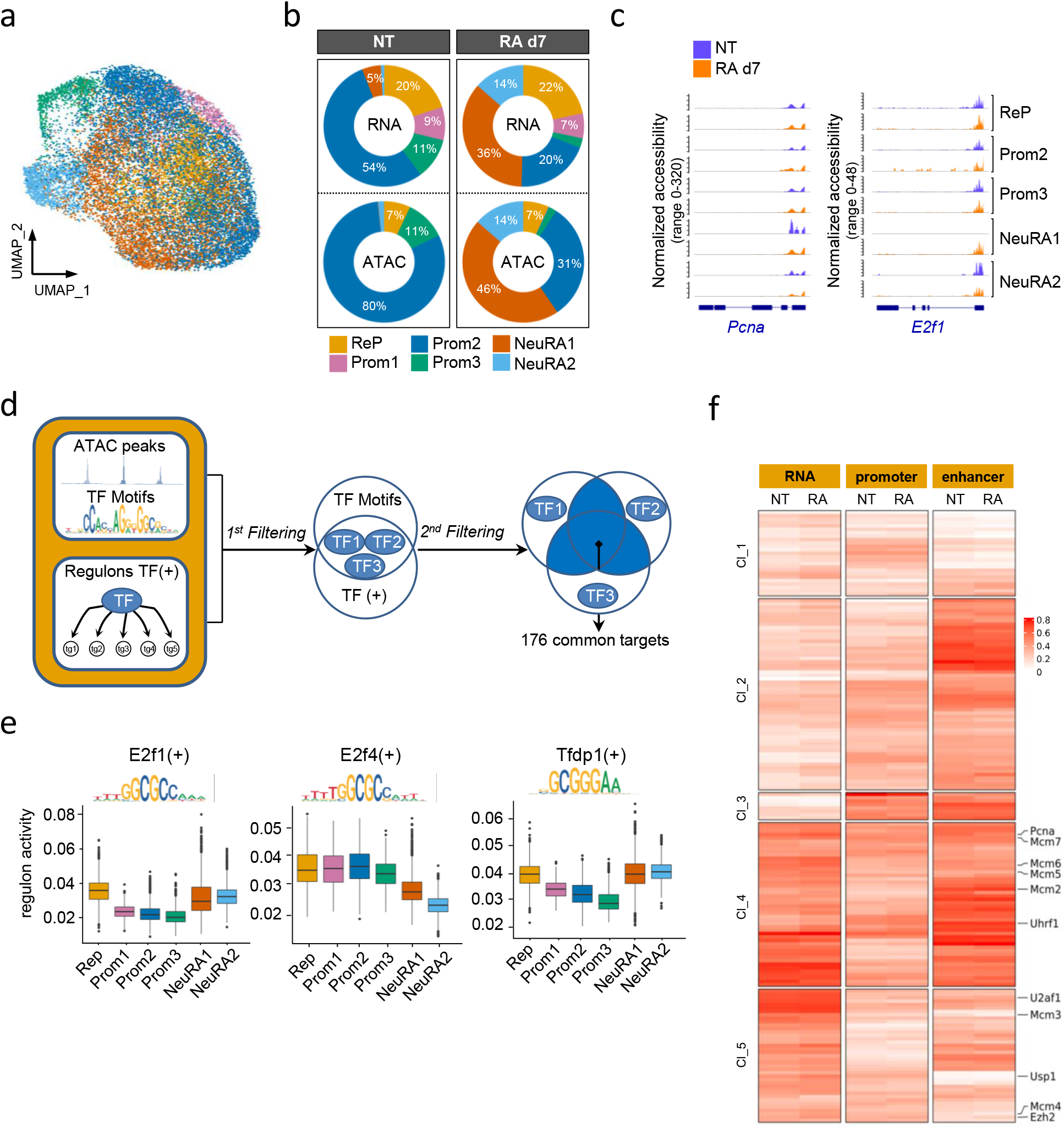
Single-cell integrative multi-omics analysis highlights chromatin genes as responsible for RA resistance in PLZF-RARA expressing cells. (**A**) UMAP visualization of integrated scRNA-seq (11,900 cells) and scATAC-seq (7,367 cells) dataset colored by cluster. (**B**) Donut plots showing the distribution of scRNA- and scATAC-seq cells in each cluster for NT and RA d7 conditions. (**C**) Coverage Plot of ATAC signal per condition in each cluster at *Pcna* and *E2f1* promoter regions. (**D**) Computational scheme to identify key regulon targets in the ReP cluster. TF motif accessibility score is calculated with Signac ^23^. TF motif markers are identified for each cluster (Table S2A). Regulon activity is scored using pySCENIC and regulon markers are identified for each cluster (Table S2B). TFs of interest are filtered by intersecting TF motifs and regulon markers in the ReP cluster (1^st^ filtering). After this filtering, 3 TFs remain. Target genes shared at least by 2 TFs are taken into account for further filtering (2^nd^ filtering). Target genes which are kept are: (i) found in all pySCENIC run, (ii) linked with positive regulons and (iii) filtered based on the sum of normalized importance (> 0.35). 176 genes are conserved (Table S3). (**E**) Boxplots showing E2f1, E2f4 and Tfdp1 (TFs obtained after the 1^st^ filtering) regulon activity in each cluster. (**F**) Heatmap showing the mRNA expression (left), the promoter accessibility (middle, ± 3kb from the TSS) and the enhancer accessibility (right, ± 50kb from the TSS minus the ± 3kb promoter region) of the 176 target genes in the ReP cluster (obtained after the 2^nd^ Filtering). Results are expressed as normalized log (mean gene activity). Hierarchical clustering is done according to the NT dataset.

Beyond cell identity, scRNA-seq and scATAC-seq data are uniquely valuable to define gene regulatory networks at the cellular scale. To decipher specific TF activity that might be associated with RA resistance, we inferred information from ATAC-seq and RNA-seq data in the ReP cells (**Figures 2D-F**). We first selected ReP TF activity by considering, in our scATAC-seq data, the accessibility of DNA motifs of TFs in the ReP cluster using the Signac chromVAR package ^23^ (**Figure S2B, Table S2A**). Selected ReP TFs were cross-referenced with master transcriptional regulators identified from scRNA-seq data using Single-Cell Regulatory Network Inference and Clustering (SCENIC) ^25^ (**Figure 2D, Table S2B**). Doing so and in line with our previous annotation, we identified 3 TFs, linked to E2F (E2F1, E2F4, TFDP1) with high transcriptional activity in the ReP cluster (**Figures 2E, S2C)**. Next, we focused on their 176 shared targets and looked at both the accessibility at gene promoters (±3 kb from the transcriptional start site [TSS]) and distal regulatory regions (enhancers) as well as at the expression levels of the target genes in the ReP cluster (**Figures 2D, 2F**). K-means clustering taking into account the three levels of information highlighted five clusters of genes (**Figure 2F, Table S3**), which could account for the heterogeneity in gene regulation inside the regulons. We observed that for most of the heatmap clusters, the average ATAC signal appeared stronger at enhancer regions than at promoters. Yet, we evidenced a dynamic in chromatin and transcriptional events starting with high chromatin opening and low RNA expression (Cl_2 and Cl_3) and ending with high RNA expression and low chromatin opening (Cl_5), in agreement with a previous observation that cis-regulatory regions were often opened prior to any gene expression ^26^. This means a sequential coordination for a given TF to regulate its targets. Two clusters of genes (Cl_4 and Cl_5) were highly expressed in comparison to the others with no impact of RA treatment on them (**Figure 2F)**. These genes were not highly expressed but modified by RA treatment in the RA-responsive NeuRA2 cells (**Figure S2D, Table S3**) suggesting a role of these genes in RA resistance. In line with our previous analysis, these highly expressed genes (*Pcna, Mcm2-7 and Ezh2*) were related to DNA repair, replication and chromatin.

The results show that RA transcriptional resistance is dependent on E2F transcriptional activity and relies on chromatin accessibility differences mainly at enhancer regions. We also pointed out heterogeneity in terms of target regulation inside of a regulon.

### EZH2 is necessary for PLZF-RARA transformation activity

EZH2 is a component of PRC2 whose canonical enzymatic activity induces the repressive histone mark H3K27me3, frequently involved in AML ^27^. To ascertain the functional relevance of EZH2 in PLZF-RARA APL cells, we first confirmed its high expression in the ReP cluster and the persistence of its expression after RA-treatment (**Figures 3A**). We also confirmed the chromatin accessibility in its promoter region at the E2F1, E2F4 and TFDP1 binding motifs (**Figure 3B**). Next, we analyzed the clonogenic activity of PLZF-RARA in the absence of EZH2 *ex vivo* by transducing BM lineage negative cells of a conditional KO *Ezh2* mouse model ^28^ with a *PLZF-RARA-IRES-GFP* retroviral construct and performing replating assay (**Figure 3C**). Consistent with our single cell data, PLZF-RARA transduction in lineage-negative cells induced an overall increase in both EZH2 and H3K27me3 levels (**Figure 3D**) while sustaining repeated replatings (**Figure S3A**). *Ezh2* deletion (Δ/Δ) induced by adding 4-OHT into the methylcellulose, even in the absence of RA, reduced the cell number and dramatically altered the replating capacity of the PLZF-RARA expressing cells (**Figure 3E**) and promoted their terminal differentiation (**Figure 3F**). *Ezh2* deletion and the consecutive H3K27me3 loss were associated with a PLZF-RARA decrease (**Figure 3D**) which was consistent with the altered replating capacity and differentiation status of the cells. This loss in replating capacity was also observed when *Ezh2* deletion was achieved *in vivo* before PLZF-RARA transduction (**Figure S3B)** or *ex vivo* after the transformation process at the second or third round of plating (**Figure S3C**). These assays suggest that Ezh2 is required for the initiation and maintenance of PLZF-RARA clonogenic activity.

**Figure 3:**
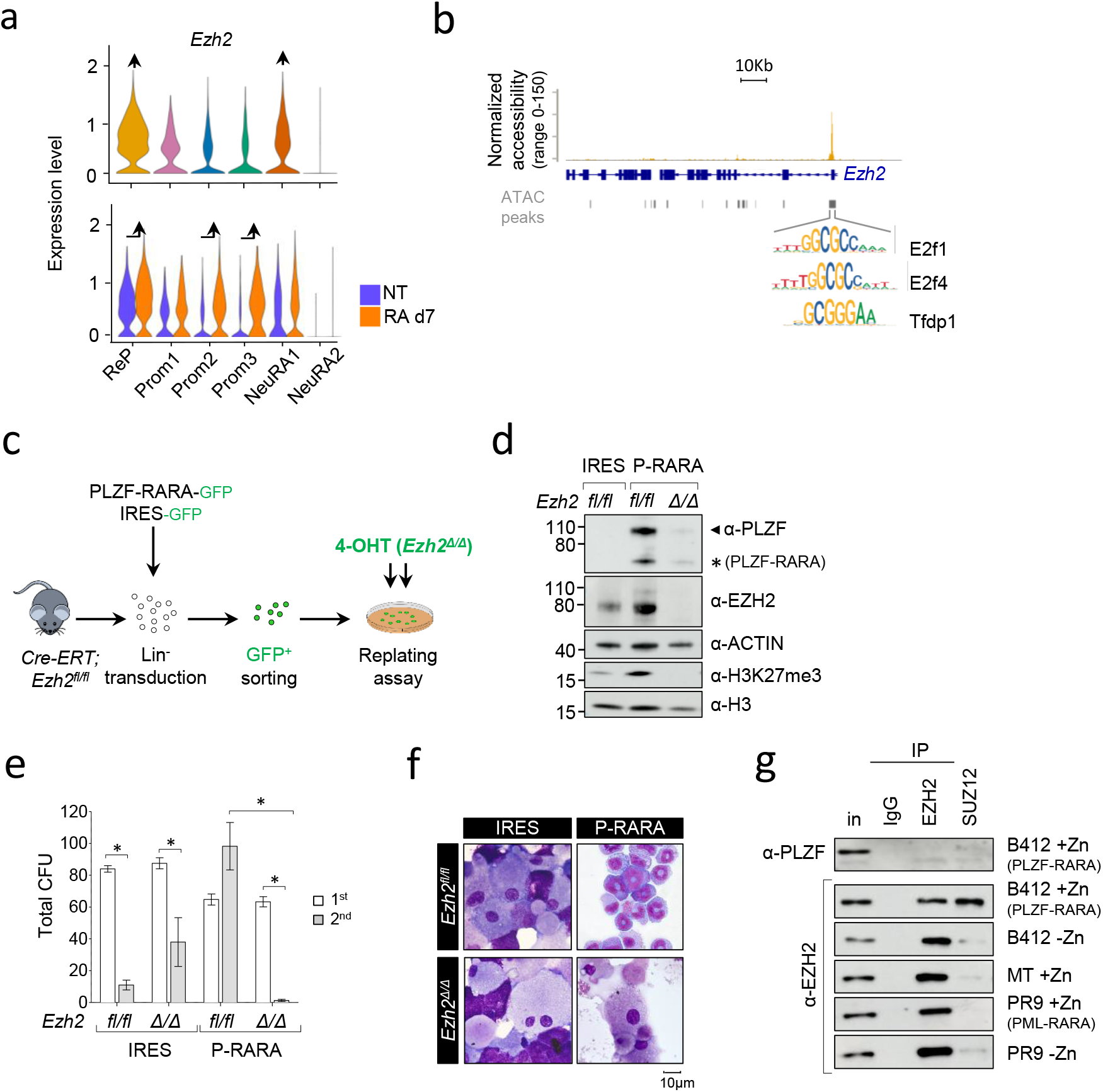
EZH2 relevance in PLZF-RARA APL. (**A**) Violin Plot showing *Ezh2* expression per cluster (upper panel) and in NT or RA treated cell conditions per cluster (lower panel). The black arrow pointing up indicate significant higher expression in the cluster against all the other clusters (average log2|FC| > 0.25 and p-value adjusted < 0.05). (**B**) Coverage Plot of ATAC signal on *Ezh2* gene in the ReP cluster. E2f1, E2f4 and Tfdp1 motifs are detected under the highlighted peak. (**C**) Experimental scheme to ascertain whether Ezh2 activity is required for PLZF-RARA transformation. Lineage negative (Lin^-^) cells purified from *Cre-ERT;Ezh2*^*fl/fl*^ are purified and transduced with an *empty-IRES-GFP* (IRES) or a *PLZF-RARA-IRES-GFP* (P-RARA) retroviral construct. Lin^-^ GFP positive cells are purified by FACS and *Ezh2* deletion in Lin-GFP positive cells is obtained by adding 150nM 4-OHT in the methylcellulose. (**D**) Global levels of PLZF-RARA (black arrow: full length, black star: degraded form), Ezh2 and H3K27me3 detected by western blotting after 4-OHT-induced Ezh2 deletion in the 2^nd^ round of plating. Actin and H3 are used as loading controls. (**E**) Replating efficiency is monitored by counting the total Colony Forming Units (CFU) of non-transformed (IRES) and PLZF-RARA transformed (P-RARA) cells in presence (*fl/fl*) or absence (*Δ/Δ)* of *Ezh2*. Results are expressed as a mean ±SD of three experiments (n=3). *p-value < 0.05. (**F**) Cell morphology of P-RARA or IRES-transduced cells in presence (*fl/fl*) or absence (*Δ/Δ)* of *Ezh2*. Representative colonies of indicated conditions after 2 rounds of plating. Cells are cytospun and observed after May-Grünwald Giemsa (MGG) staining; magnification 64X; bar 10 µm. (**G**) Nuclear extracts of U937 cells treated or not with ZnSO4 (Zn) immunoprecipitated with anti-IgG, anti-EZH2 or anti-SUZ12 antibodies. IPs are immunoblotted with anti-PLZF (upper panel) or anti-EZH2 antibodies (lower panel). Inputs (in) represent 2% of samples processed in each IP. U937 B412: PLZF-RARA Zn-inducible; U937 MT: parental cells; U937 PR9: PML-RARA Zn-inducible.

We next investigated a potential interaction between the oncogenic fusion protein PLZF-RARA and EZH2 using a myeloid cell line (U937-B412) with zinc inducible PLZF-RARA expression (**Figure 3G**) and HEK293T cells overexpressing a Flag-tagged EZH2 and PLZF-RARA (**Figure S3D**). In both systems, we could not demonstrate any interaction between PLZF-RARA and EZH2 or SUZ12, another component of PRC2 (**Figure 3G; Figure S3D**). However, when probing our SUZ12 IP with an anti-EZH2 antibody, we evidenced a stronger interaction between EZH2 and SUZ12 in the presence of PLZF-RARA than without (**Figure 3G)**. The increase in interaction was not found in the presence of PML-RARA, demonstrating a specificity of the stabilizing effect of PLZF-RARA on the EZH2/SUZ12 complex (**Figure 3G**).

Collectively, these results reveal a dependency of PLZF-RARA expressing leukemic cells upon EZH2 whose activity may itself be modified by the presence of the fusion protein.

### PLZF-RARA induced H3K27me3 level at specific enhancers genes that marks RA relapse initiating cells

Next, we investigated the effect of PLZF-RARA expression on EZH2 chromatin activity. We compared the epigenetic landscape of PLZF-RARA promyelocytes with the Granulocyte-Monocyte-Progenitor (GMP) compartment, which is the closest cell compartment to promyelocytes according to its transcriptional signature (**Figure S4A**). Because our scATAC-seq data revealed stronger chromatin opening at enhancer than promoter regions (**Figure 2F**) and since cis-regulatory enhancer elements are known to influence the development of leukemia ^29^, we mapped the four histone marks (H3K27ac, H3K27me3, H3K4me1 and H3K4me3) that allow to discriminate active (H3K27ac-enriched) and poised (H3K27me3-enriched) enhancers ^30^ (**Figure S4B**). By comparing PLZF-RARA promyelocytes with GMP genomic enhancer distribution, we found that PLZF-RARA expression significantly modified poised (H3K27me3-enriched) enhancer positions (68% of them were specific to PLZF-RARA condition) while minimally affecting the active enhancers (70 % of them were overlapping in the two conditions) (**Figure 4A**). These new H3K27me3-enriched enhancer regions in PLZF-RARA condition reflected a shift in H3K27me3 enrichment from enhancers regulating developmental processes to those regulating kinase signaling and bloodstream systems (**Figure S4C**). Next, we focused on switched enhancers which were marked by H3K27ac in GMP condition but gained H3K27me3 with PLZF-RARA expression (**Figures 4A, S4D**). Although these switched enhancers were in the minority (175 of a total of 3876 PLZF-RARA induced poised enhancers), they exerted a specific effect on the expression of their nearby genes in the ReP cluster, in which the expression was lower compared with the other clusters (**Figure 4B, upper panel**). In contrast, *de novo* enhancer-associated genes were equally weakly expressed in all scRNA-seq clusters (**Figure 4B, lower panel**). Gene Ontology (GO) analysis revealed that switched enhancer-related genes were associated with myeloid cell differentiation and pro-apoptotic program regulatory terms (**Figure 4C**). In addition, their expression was decreased in the presence of PLZF-RARA (**Figure S4E**). This indicates that PLZF-RARA expression resulted in a redistribution of repressed enhancers whose activity limits apoptosis and myeloid differentiation and suggests that the change in the specific poised enhancer switch of the ReP cluster cells may be related to the lack of response to RA.

**Figure 4:**
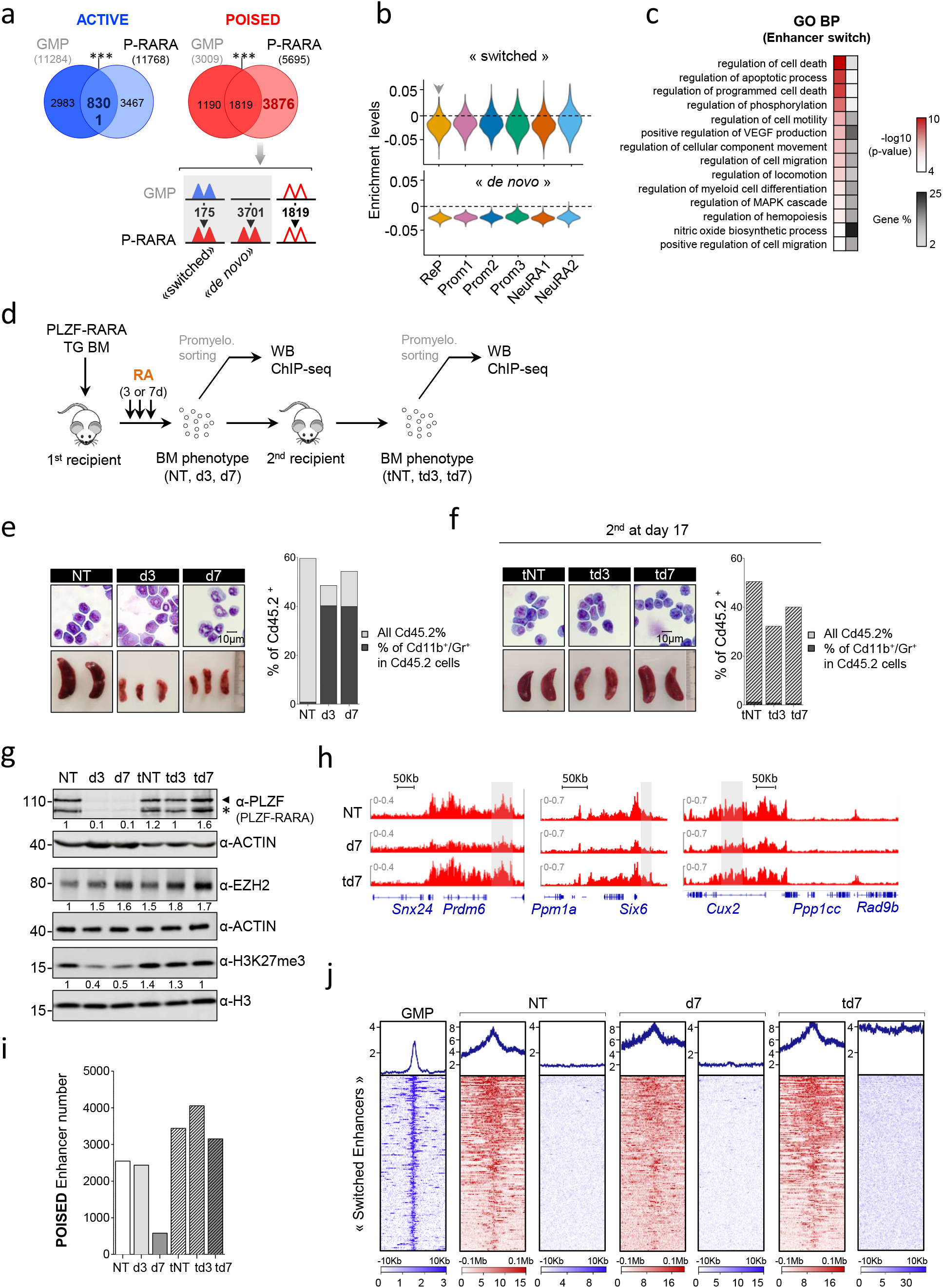
PLZF-RARA induced H3K27me3 level at specific enhancers genes that marks RA relapse initiating cells. (**A**) Enhancer distribution in PLZF-RARA promyelocytes is compared to those observed in normal GMPs (Granulocyte-Monocyte Progenitors) (GSE124190). The Venn diagrams show the overlap of Active (left panel) and Poised enhancers (right panel) in GMP and in PLZF-RARA (P-RARA) conditions. *** p-value < 0.001. The poised enhancer dynamics upon PLZF-RARA expression is schematized below. Triangles represent enhancers, colors indicate their activity; active are colored in blue, poised in red. Numbers indicate the enhancers that are found in PLZF-RARA condition. When no changes in enhancer activity is observed between the two conditions, triangles are empty. (**B**) Violin plots showing “ switched “ (upper panel) and “ *de novo* Poised” (lower panel) signature scores per cluster. Signature score represents the global expression of annotated genes for the selected signature identified by scRNA-seq. The gray arrow pointing down indicates significant lower expression in the cluster against all the others (average score difference > 0.005 and p-value adjusted < 0.05). (**C**) Gene Ontology (GO) analysis using GREAT tools of enhancer nearby genes. Red scale values indicate the p-value (-log10) and gray scale values illustrate gene % (*i*.*e* number of genes observed/total number of genes within each GO term). (**D**) Experimental scheme to assess chromatin events associated with retinoic acid (RA) resistance. PLZF-RARA TG bone marrow is transplanted into recipient mice. Ten days after, mice are injected with corn oil (NT) or with RA for 3 or 7 days (d3 and d7) and sacrificed. Treated BMs are immunophenotyped and reinjected into new recipient mice (tNT, td3, td7). Secondary transplanted mice are not treated (Ø) and sacrificed at day 17 post transplantation. Promyelocytes are FACS sorted at each sacrifice for western blotting (WB) and epigenomic analyses. (**E-F**) Leukemia evolution analyzed by cell morphology with May Grunwald Giemsa (MGG) staining (left upper panel); magnification 64X; and spleen size (bar in centimeter) (left lower panel) and FACS analysis (Cd45.2, Cd11b and Gr1 markers are monitored) (right panel). In (**E**) impact of 3 or 7 days of RA treatment on PLZF-RARA leukemia (NT, d3, d7) and in (**F**) leukemia relapse evaluation of transplanted untreated (tNT) and RA-treated BM (td3, td7) at day 17 post transplantation. (**G**) Global levels of PLZF-RARA (black arrow: full length, black star: degraded form), EZH2 and H3K27me3 detected by western blotting (WB). Actin and H3 are used as loading controls. Signal intensity is measured and normalized according to the loading control and to the untreated condition (NT). (**H**) Representative Integrative Genomics Viewer (IGV) tracks of H3K27me3 in NT, d7 and td7 conditions. The gray box underlines enhancer coordinates. (**I**) Total number of poised enhancers in each condition. (**J**) Plot Heatmap of H3K27ac and H3K27me3 signals in GMP, NT, d7 and td7 conditions at switched enhancers (Figure 4A, 175 enhancers). H3K27ac and H3K27me3 signals are plotted 10Kb and 0.1 Mb upstream and downstream the enhancer center.

To determine the role of EZH2 in RA resistance, we investigated the effect of RA on EZH2 chromatin activity after sequential transplantations (**Figure 4D**). As previously observed ^13^, RA treatment, although inducing a clear differentiation of blasts (**Figure 4E)**, did not eradicate relapse-initiating cells, since relapse was observed at day 17 post-transplant in mice transplanted with RA-treated BMs (**Figure 4F**). We showed that RA had a contrasting effect on EZH2 in leukemic cells. While increasing its protein level, which was consistent with our scRNA-seq data (**Figure 3A**), it decreased the overall level of H3K27me3 as early as day 3 of treatment (**Figure 4G**). Interestingly, leukemia relapse of RA-treated BM was characterized with a restoration of high levels of H3K27me3 in the leukemic cells (**Figure 4G**). This suggests that RA-induced differentiated cells maintained EZH2 expression but not its methyl transferase activity. More precisely, RA decreased H3K27me3 signal at poised enhancers (**Figure 4H**) resulting in a diminution in numbers of these enhancers after 7 days of treatment (**Figure 4I**). Clearly, leukemia relapse was characterized by both the restoration of high levels of H3K27me3 at the enhancer regions and the high number of H3K27me3-enriched enhancers (**Figures 4H-I**). Because PLZF-RARA cells respond differently to RA and the ReP cluster may be responsible for RA resistance, we measured the levels of H3K27me3/H3K27ac after RA treatment and at relapse at the 175 switched enhancers (described in **Figure 4A**). We found that neither RA treatment (3 or 7 days) nor the following transplantations erased the H3K27me3 signal at these enhancers, for which H3K27ac stayed lower than in GMP (**Figures 4J & S5A**). Consistently, the signature of the switched enhancers was not impacted by RA in our scRNA-seq data (**Figure S5B**).

Collectively, these results show that PLZF-RARA modifies EZH2 activity by redirecting the H3K27me3 on a subset of enhancers which are not affected by RA treatment. We also highlight a discrepancy between EZH2 protein level and its methyltransferase activity during RA-induced differentiation.

### Elimination of EZH2 eradicates relapse leukemia initiating cells

Because EZH2 is necessary for PLZF-RARA induced transformation and high H3K27me3 level at specific loci is a marker of resistance, we questioned the pertinence of targeting EZH2 with GSK126 in combination with RA to overcome resistance. To test this hypothesis, we treated PLZF-RARA TG transplanted mice with GSK126 and/or with RA and transplanted the treated BMs into new recipients to follow disease progression according to the pre-treatment (**Figure 5A**). At first, the inhibitory effect of GSK on H3K27me3 level in the treated BM, which was more pronounced when the drug was administered in combination with RA, was confirmed (**Figure 5B**). However, GSK treatment did not affect the % of PLZF-RARA TG cells as reflected by the % of Cd45.2 positive cells in total BM nor, in contrast to RA, impacted on disease progression and blast differentiation (**Figure 5C**). Besides, no synergistic effect of GSK with RA was observed (**Figure 5C**). Leukemia relapse was monitored by following the transplantation of the treated BMs in new recipient mice (**Figure 5D**). At day 13 post transplantation, GSK treated BM expanded as untreated BM suggesting that GSK alone did not impact the relapse initiating cells. A delay in leukemia progression was observed in the RA-treated BM, but addition of the GSK did not change in any instance the leukemia progression. To rule out accessibility and dosage problems that could be faced using animal models, we compared the effect of GSK treatment alone or in combination with RA on PLZF-RARA replating capacity (**Figure S6A**). As for the *in vivo* experiment, GSK alone or in combination with RA had no effect on the replating capacity of PLZF-RARA *ex vivo* (**Figure S6B)** although inducing some cell differentiation features (**Figure S6C**). These data showed that despite a strong dependency of PLZF-RARA cell transformation on EZH2, inhibition of its catalytic activity is not sufficient to promote APL clearance and further suggested that PLZF-RARA APL depends on a non-canonical activity of EZH2. To explore this possibility, we took advantage of a new commercially available EZH2 degrader, MS1943 (MS) ^31^. PLZF-RARA TG BM was treated with MS and re-transplanted in new recipient mice (**Figure 5E)**. We observed a strong effect of the MS on cell viability (**Figure 5F**) concomitant to its ability to erase H3K27me3 level and efficiently degraded EZH2 (**Figure S6D)**. This effect was greater in cells expressing PLZF-RARA than in those not expressing it (**Figure S6E**). Reinjecting alive MS-treated PLZF-RARA BM cells in recipient mice significantly prolonged the survival of the mice, confirming the importance of EZH2 non-canonical activity in PLZF-RARA leukemia development (**Figure 5G**). This gain of survival upon MS treatment was associated with a strong and specific decrease in the expression of RA resistant ReP cluster markers (e-g: E2f1, Rad51ap1, Pcna, Mcm2 or Uhrf1), which were less affected with the GSK treatment (**Figures 5H**).

**Figure 5:**
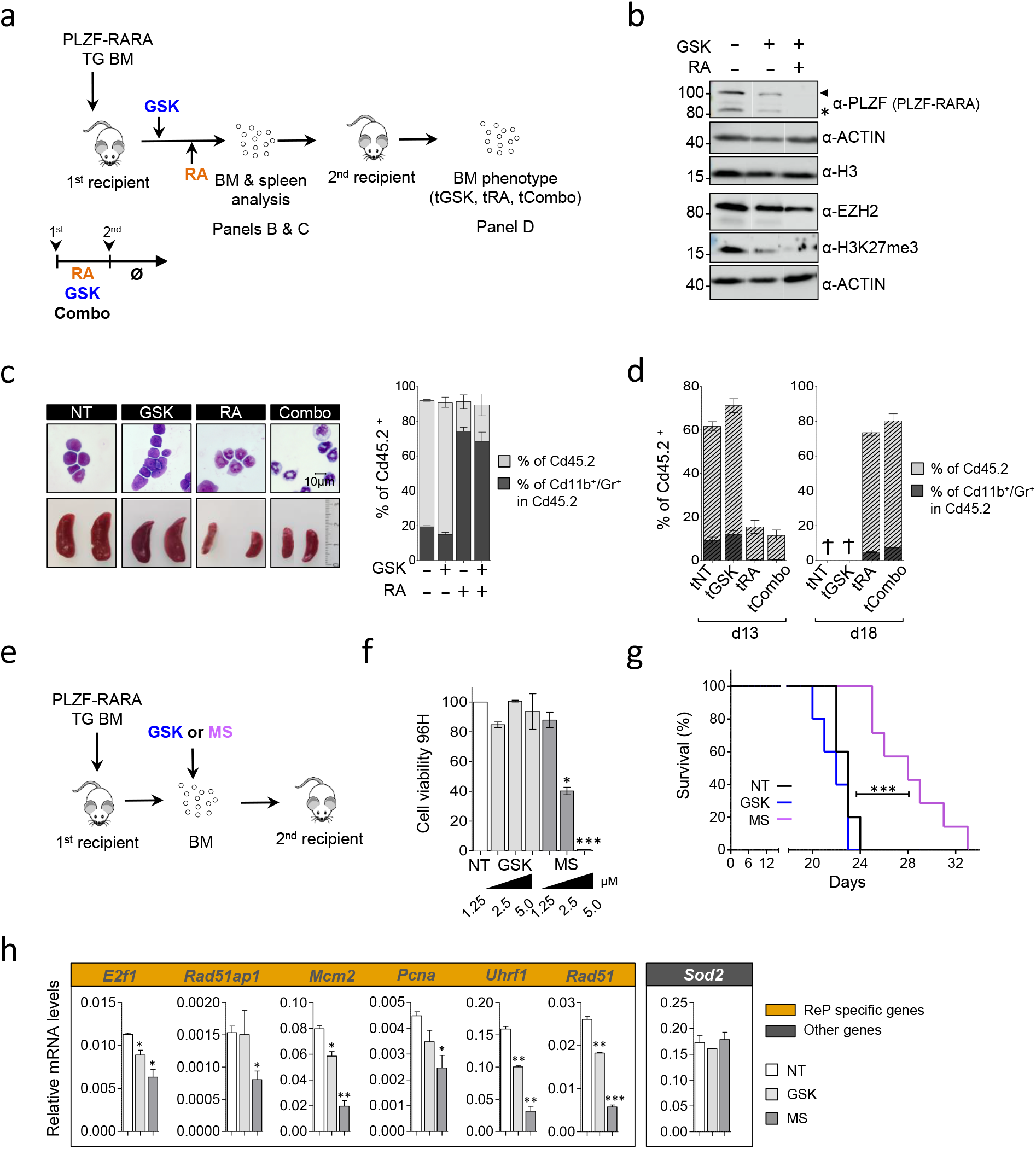
Impact of targeting EZH2 on APL Progression. **(A)** Experimental scheme of the analysis of RA, GSK or combo treated bone marrow. PLZF-RARA TG bone marrow is transplanted into recipient mice. Seven days after, mice are injected for 3 days with GSK126 (GSK) or with corn oil (NT) followed by 7 consecutive days of RA (RA). After treatments (NT: corn oil, GSK, RA, combo: GSK and RA) mice are sacrificed and treated-BM are immunophenotyped and reinjected into new recipient mice (tNT, tGSK, tRA, tCombo). At this time treatments are stopped (Ø). Mice are sacrificed 15-20 days after the secondary transplantation. (**B**) Global levels of PLZF-RARA (black arrow: full length, black star: degraded form), Ezh2 and H3K27me3 detected by western blotting (WB). Actin and H3 are used as loading controls. (**C**) Impact of GSK, RA and combo treatments on PLZF-RARA leukemia (NT, GSK, RA, Combo). Cell morphology analyzed by May Grünwald Giemsa (MGG) staining (left upper panel); magnification 64X; bar 10 µm; spleen size (bar in centimeter) (left lower panel) and FACS analysis (Cd45.2, Cd11b and Gr1 markers are monitored) (right panel). (**D**) Evaluation of leukemia relapse monitored by FACS analysis (Cd45.2, Cd11b and Gr1 markers are monitored) at day 13 (d13) and day 18 (d18) after the secondary transplantation (tNT, tGSK, tRA, tCombo). (**E**) Scheme resuming the protocol to evaluate the effect of MS treatment on PLZF-RARA BM. (**F**) Cell viability of PLZF-RARA TG cells upon GSK126 (GSK) or MS1943 (MS) treatments monitored by bioluminescence. Results are expressed by percent of living cells and normalized to the untreated condition (NT). results are expressed as the mean ±SD of three independent experiments (n=3). *p-value < 0.05, *** p-value < 0.001. (**G**) Survival rate (Kaplan Meier survival analysis, https://www.statskingdom.com/kaplan-meier.html) of mice transplanted with untreated (NT, black curve), MS1943 (MS, 2.5µM, purple curve) or GSK126 (GSK, 2.5µM, blue curve) pretreated PLZF-RARA TG bone marrow. *** p-value < 0.001. (**H**) mRNA levels of selected genes in the ReP signature monitored by qPCR. Sod2 is presented as a control gene not affected by RA. mRNA values are normalized to *β2microglobulin* and are expressed as a mean SD of two independent experiments (n=4). *p-value < 0.05, *** p-value < 0.001.

Altogether, these results demonstrate that targeting EZH2 methyltransferase activity is not sufficient to eradicate relapse leukemia initiating cells and suggest elimination of EZH2 to overcome RA resistance.

## DISCUSSION

Cancer cell heterogeneity is a major driver of therapy resistance and disease progression. To address this problematic, we focused on the effect of RA treatment in APL cells expressing PLZF-RARA, a paradigm to study oncogenesis and RA resistance. Given the unique differentiating properties of RA, many studies are being evaluated for treatment of non-APL AML ^32^ and solid tumors ^33^, and a better understanding of resistance mechanisms could widen its use. We demonstrate the power of single-cell multi-omics to understand cancer cell heterogeneity and its impact on treatment resistance. By leveraging our own single-cell data, we reveal a novel mechanism of resistance, involving the epigenetic enzyme EZH2, opening up a therapeutic opportunity (**Figure 6**).

**Figure 6:**
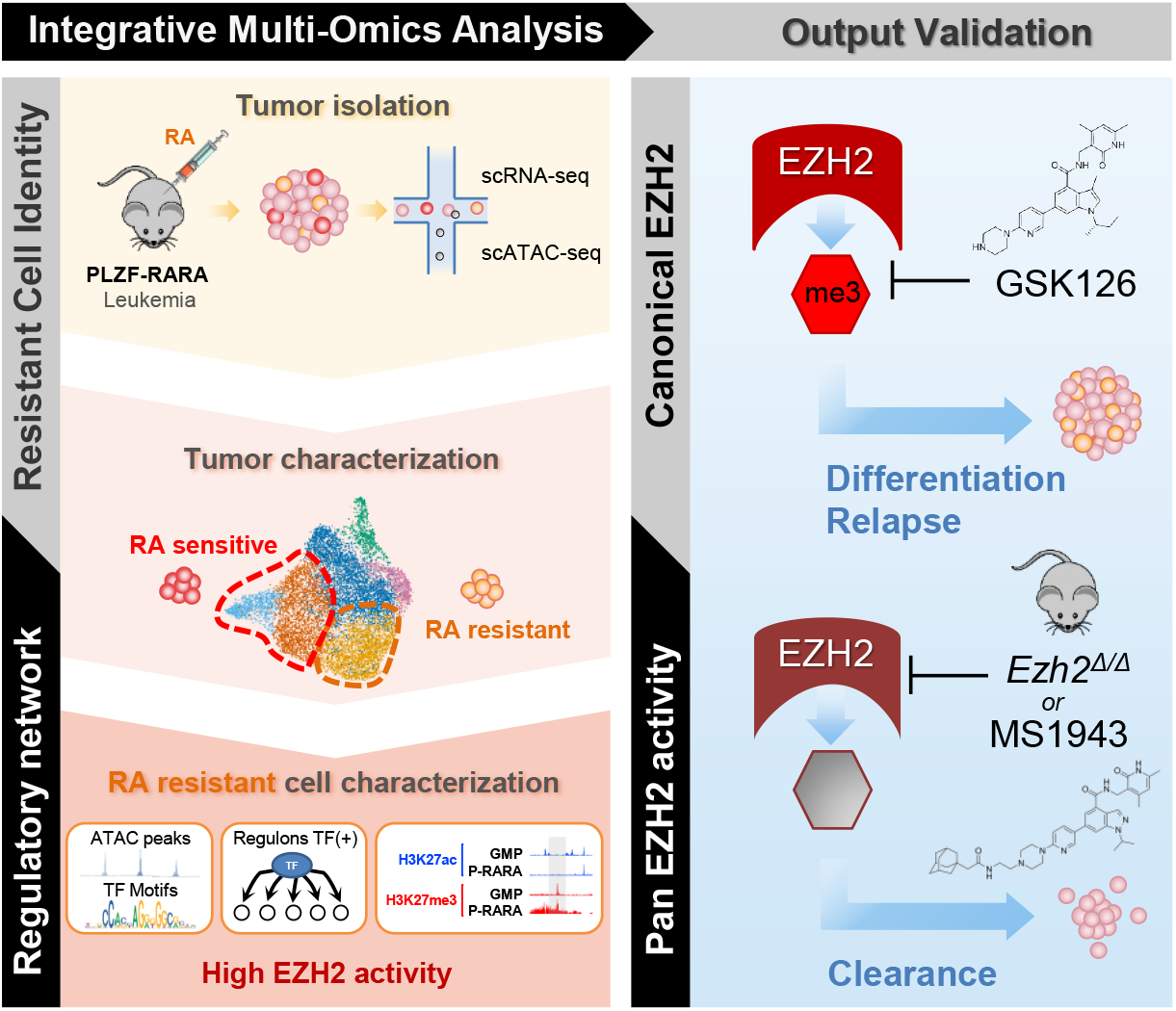
Summary abstract.

A notable aspect of our findings is the heterogeneity of PLZF-RARA transformed cells with their different responses to RA. Clonal diversity and evolution patterns of AML by high-throughput single-cell genomics (scRNA-seq and scDNA-seq) have revealed hierarchies in the AML with heterogeneity correlating with their underlying genetic alterations ^34 35^. Here, we studied a unique gene alteration and showed that in the presence of a single oncogenic event, cells are not homogeneously transformed and respond differently to RA. We identified different subgroups of transformed promyelocytes, based on differential transcriptional activity, which reflect the APL hierarchy as well as spontaneous differentiation. By probing the heterogeneity of the leukemic bulk before and after RA treatment, we identified and characterized a group of RA resistant cells, which we named ReP. ReP cells remained transcriptionally and at the chromatin level unperturbed by RA treatment and kept their leukemic activity. What could make these cells the RA-resistant cells? Strikingly, this group of ReP cells differed from the other groups by a weak myeloid differentiation program and a marked replication/repair program. In accordance with these features, enrichment analysis highlighted a high E2F activity, which could sustain the resistance of the cells considering its role in promoting leukemic cell survival ^36^, apoptosis evasion ^37^ and resistance to cisplatin treatment ^38^. In addition, we showed that this group of cells maintains PLZF-RARA expression after RA treatment, which is known to act as a competitive transcriptional repressor of RARA ^15^. Although we cannot determine whether the maintenance of PLZF-RARA is the cause or consequence of RA unresponsiveness, it underscores the dependence of the leukemia on the driver oncogene.

Our scATAC-seq single experiment allowed us to study E2F activity in the RA resistant cluster by analyzing its targets, which are known to mediate its oncogenic activity ^39^. Among the E2F targets, we identified EZH2 as an interesting epigenetic vulnerability to overcome RA resistance. EZH2 deregulations are related to cancer initiation, metastasis, immunity, metabolism, and drug resistance in a wide variety of cancers ^40^. However, targeting EZH2 in leukemia may be complex due to its tissue and action mode specificity in the hematopoietic compartment ^28^. This complexity is also observed within AMLs: EZH2 exerts an oncogenic function by enhancing differentiation blockade during AML maintenance ^41^, while it acts as a tumor suppressor during leukemia initiation ^18^. Here, we showed that survival of PLZF-RARA transformed cells is dependent on EZH2 activity. By integrating RNA single cell data with ChIP-seq data, we demonstrated that EZH2 methyltransferase is redirected to apoptotic genes by PLZF-RARA and this redirection has transcriptional consequences in the ReP cells that resist to RA treatment. Interestingly, we could not evidence a direct interaction between EZH2 and PLZF-RARA as we previously did for PLZF ^42^. However, we clearly evidenced a stabilization of EZH2 and SUZ12 association upon PLZF-RARA induction. Such PRC2 stabilization may change the methyltransferase activity of EZH2 ^43^. Thus, we revealed that the oncogenic activity of PLZF-RARA depends on the integrity of EZH2 but in turn PLZF-RARA stabilizes EZH2/SUZ12, this loop of regulation could be the key target to overcome RA resistance.

Another relevant finding of our study is the little effect of inhibiting EZH2 methyltransferase activity on disease progression and relapse. Our results clearly showed a dual effect of the EZH2 methyltransferase inhibitor, having an eliminatory effect on H3K27me3 at differentiating genes, thus promoting differentiation, while maintaining high EZH2 expression. One of the plausible hypotheses is that inhibition of EZH2 catalytic activity has a differentiating effect on PLZF-RARA cells but does not affect the cells that compose the ReP cluster, which are characterized by a strong resistance signature. These cells may depend on an oncogenic but non-catalytic activity of EZH2. Many cancers have been shown to rely on an EZH2 oncogenic effect that is not solely based on its histone transferase activity. This is the case of estrogen receptor (ER)-negative basal-like breast cancers ^44^, AR dependent prostate cancers ^45^ or SWI/SNF mutant tumors that rely on both the catalytic and non-catalytic activities of EZH2. Interestingly, as for PLZF-RARA leukemic cells, GSK126 only showed limited efficacy in SWI/SNF-mutant cancer cells ^46^. This prompts us to use an EZH2-selective degrader ^31^. We showed that the use of an EZH2-selective degrader significantly reduced growth and viability of PLZF-RARA expressing leukemic cells and increased the survival of the mice transplanted with pre-treated leukemia. The development of EZH2-Based PROTACs to degrade the PRC2 complex, which targets the enzymatic and non-enzymatic activities of EZH2 ^47^ holds great promise for the future treatment of cancers dependent on the non-catalytic activity of EZH2.

In conclusion, we have validated the use of combining chromatin and expression analyses at the single cell resolution to identify treatment resistant cells and characterize their activity. We revealed a molecular mechanism of RA resistance involving EZH2, which is potential therapeutic target.

## RESOURCE AVAILABILITY

### Data availability

The raw ChIP-seq, scATAC-seq and scRNA-seq data (fastq files) are deposited under the accession number GSE181190. The repository also contains the bigwig, the peak matrix and the count matrix for the ChIP-seq, scATAC-seq and scRNA-seq respectively. All R and python codes used for data analysis are integrated in a global snakemake workflow available at https://gitcrcm.marseille.inserm.fr/herault/GMP_PRARA.

## ACKNOWLEDGMENTS

We thank Dr. Pier Pandolfi for sharing the PLZF-RARA TG mouse model; Dr Valerie. Lallemand-Breitenbach for facilitating the access to the PLZF-RARA TG bone marrow; Dr Christophe Lachaud for helpful discussion and for reading the manuscript and members of the Duprez laboratory for helpful discussions. The authors wish to thank the animal support facility of the Centre de Recherche en Cancérologie de Marseille, the flow cytometry and cell sorting platform, the CIBI and DISC platforms for computational analyses and support.

## AUTHOR CONTRIBUTIONS

M.P. and E.D. developed the original hypothesis and designed experiments. M.P. with the help from N.P., K.S., W.K., L.N., M.O. and N.C. performed experiments and/or analyzed data. A.M and L.H. carried out computational analyses of the scRNA-seq and scATAC-seq. L.H. developed the pipeline for scRNA-seq and scATAC-seq data integration. M.P. and J.V. conducted ChIP-seq analyses. A.I. and D.B. provided reagents and discussed results. M.P., E.D. and A.M. designed the figures and M.P. and E.D. wrote the manuscript. All co-authors proofread the manuscript.

