## Supplemental Figures for "Single-cell multi-omics analyses reveal EZH2 as a main driver of retinoic acid resistance in PLZF-RARA leukemia"

### Figure S1 (related to Figure 1), *legends on next page*

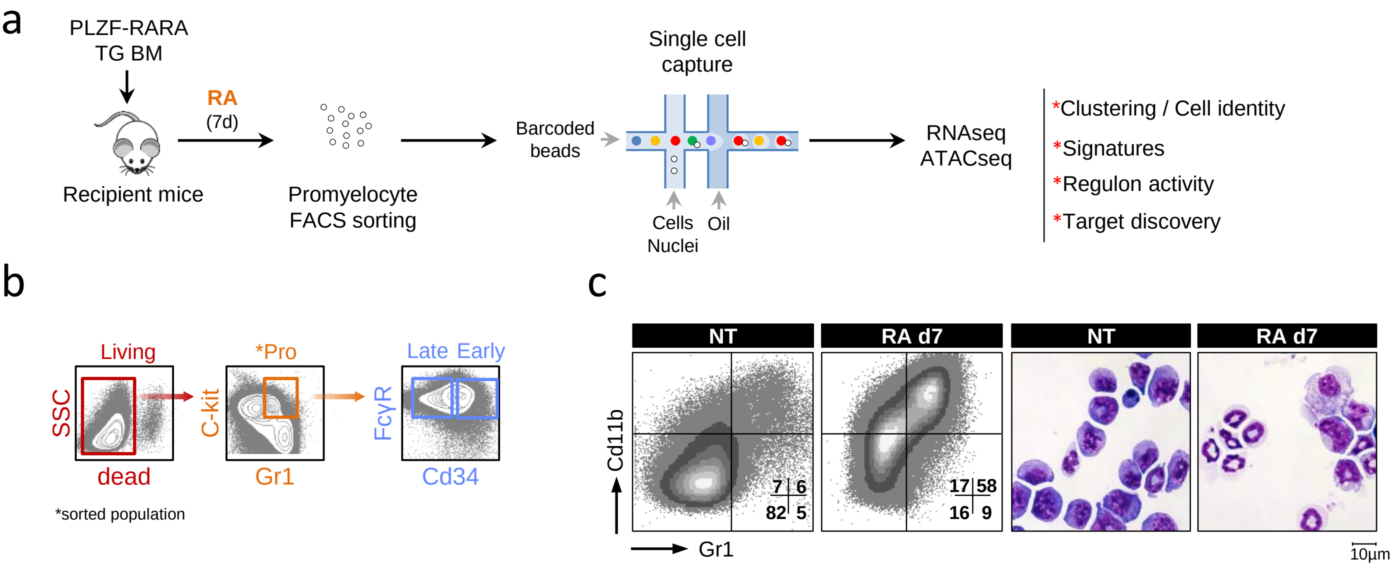


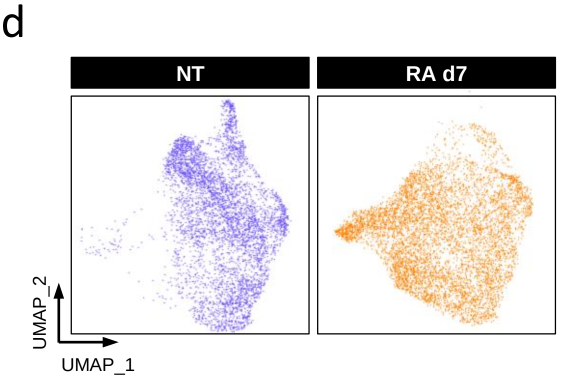


e

| Selected enrichments | p.value | Clusters |
|---|---|---|
| BP : DNA replication | 1.69e-44 | ReP |
| REA : Neutrophil degranulation | 3.73e-16 | Prom1 |
| BP : Leukocyte migration | 1.10e-06 | Prom2 |
| REA : Neutrophil degranulation | 7.77e-16 | Prom3 |
| REA : Neutrophil degranulation | 1.14e-2 | NeuRA1 |
| REA : Neutrophil degranulation | 1.77e-4 | NeuRA2 |

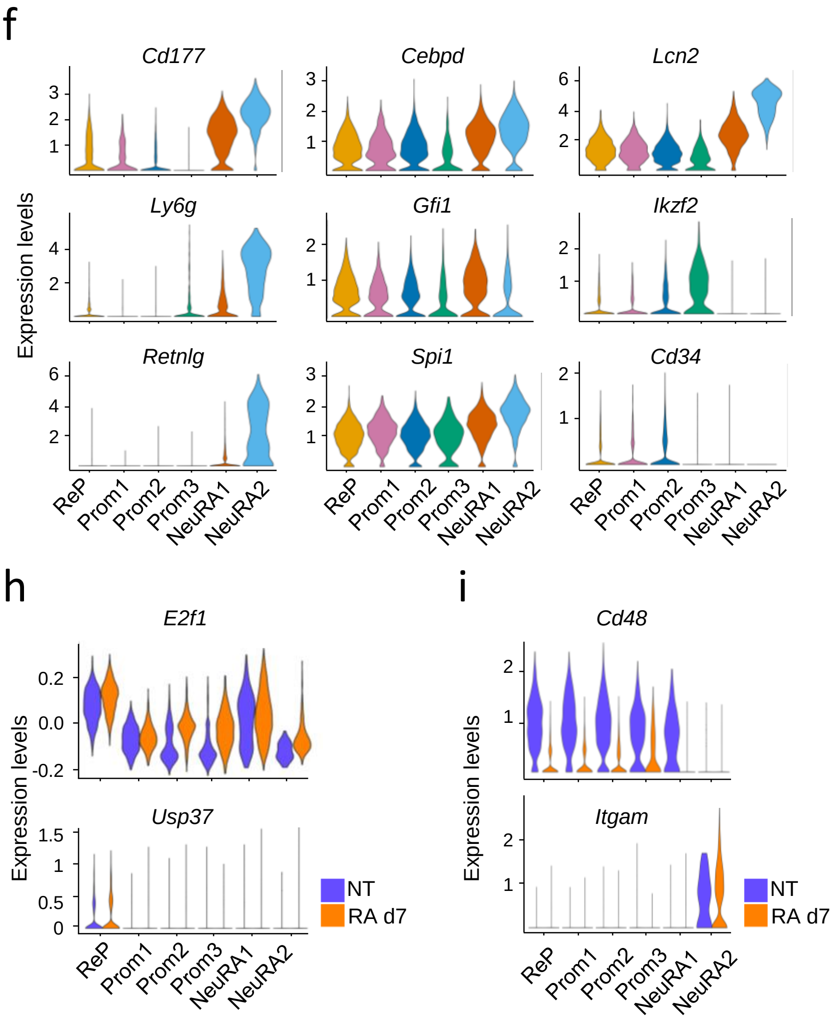

### Figure S1 (related to Figure 1): *legends*

**Figure S1 related to Figure 1:** (**A**) Experimental scheme. PLZF-RARA TG bone marrow is transplanted into recipient mice. Ten days after transplantation, mice are intraperitoneally injected with corn oil (NT: no treatment) or with 0.8-1 mg/day of the differentiating retinoic acid (RA) agent for 7 days (d7). (**B**) Gating strategy for isolating promyelocytes. C-Kit$^+$, Gr1$^+$ promyelocytes are FACS-sorted for single cell analyses. Promyelocytes are phenotypically divided into 2 sub-populations : late (C-Kit$^+$, Gr1$^+$, Cd34$^-$, FcγR$^+$) and early (C-Kit$^+$, Gr1$^+$, Cd34$^+$, FcγR$^+$) promyelocytes (**C**) RA exposure leads to promyelocyte differentiation. Left panel: FACS analysis showing an increase in double positive Cd11b and Gr1 cells upon RA. Right panel: cell morphology analyzed by May Grunwald Giemsa (MGG) staining (magnification 64X; bar 10 µm) and showing increased granulations and nuclei lobulations and diminished basophilia and nuclear/cytoplasmic ratio in RA treated promyelocytes. (**D**) UMAP visualization of NT (5,106 cells) and RA (6,794 cells) promyelocytes. (**E**) Gene set enrichment analysis of cluster markers (BP: Gene Ontology Biological Processes; REA: Reactome pathways; Tables S1A-C). (**F**) Violin plots of some selected NeuRA1 and NeuRA2 markers involved in terminal myeloid diferentiation (up/down regulated compared to the other clusters) (Table S1B) : *Cd177*, *Cebpd*, *Lcn2*, *Ly6g*, *Gfi1*, *Retnlg*, *Spi1*, *Ikzf2*, *Cd34* (average $\log_2|FC| > 0.25$ and p-value adjusted $< 0.05$). (**G**) Violin plots showing some selected ReP markers (Table S1A) : *Pcna*, *Rad51*, *Mcm3* (average $\log_2|FC| > 0.25$ and p-value adjusted $<$ 0.05). (**H**) Violin plots showing E2f signature and *Usp37* expression in NT and RA d7 conditions. (**I**) Violin plots showing *Cd48* and *Itgam* (*Cd11b*) expression in NT and RA d7 conditions per cluster.

### Figure S2 (related to Figure 2)

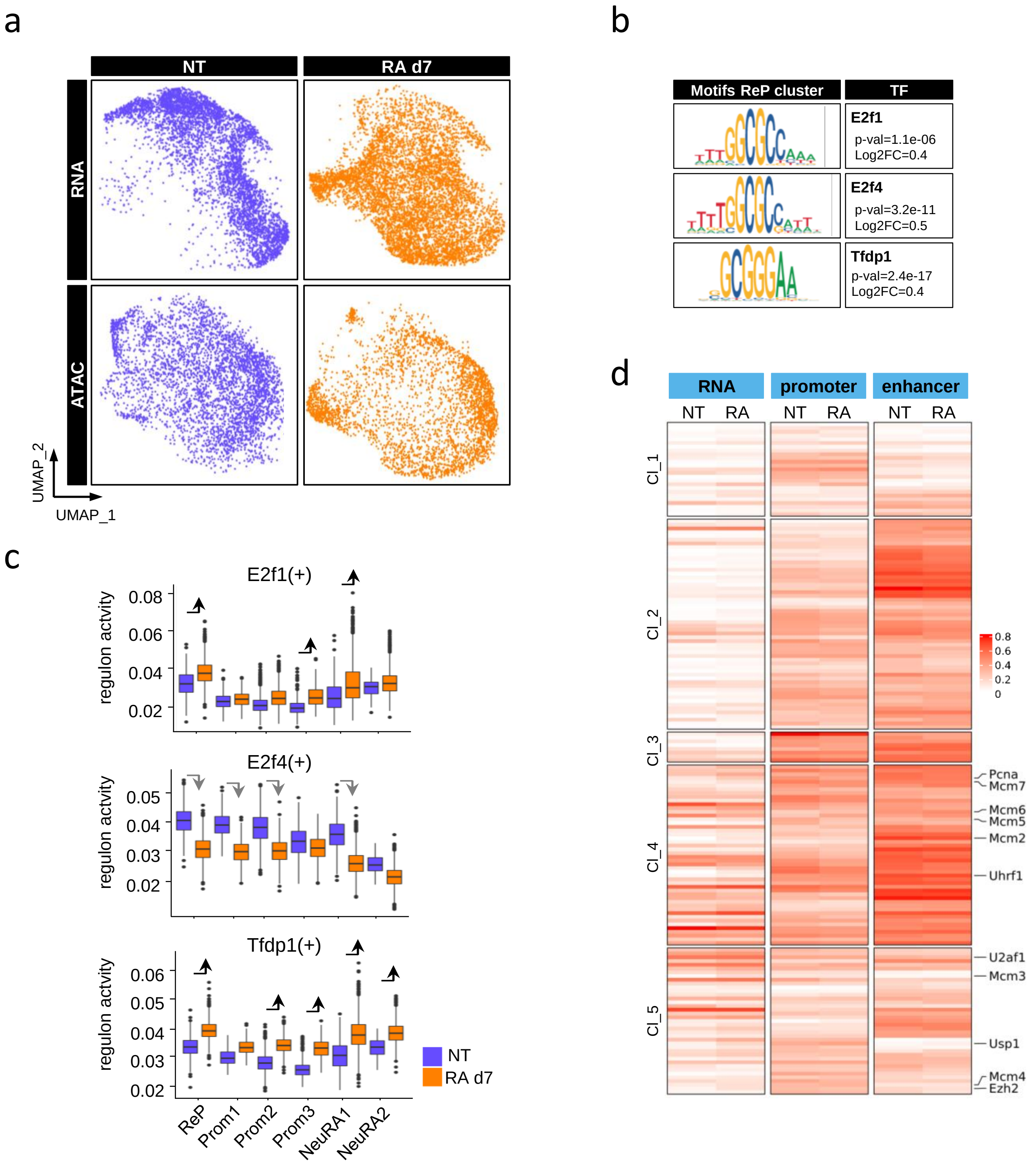


**Figure S2 related to Figure 2.** (**A**) UMAP visualization of integrated scRNA-seq and scATAC-seq from NT (RNA: 5,106 cells; ATAC: 3,954 cells) and RA d7 (RNA: 6,794 cells; ATAC: 3,413) datasets. (**B**) motifs and scores (p-value and $\log_2|FC|$) of the three marker TFs in the ReP clusters identified Figure 2D. (**C**) Boxplots showing E2f1, E2f4 and Tfdp1 (TFs obtained after the 1st filtering) regulon activity per condition in each cluster. The gray/black arrows pointing down/up indicate significant lower/higher score in one condition compared to the other (average score difference > 0.005 and p-value adjusted < 0.05). (**D**) Heatmap showing the mRNA expression (left), the promoter accessibility (middle, ± 3kb from the TSS) and the enhancer accessibility (right, ± 50kb from the TSS minus the ± 3kb promoter region) in the NeuRA2 cluster of the 176 target genes analyzed in the ReP cluster (Figure 2F). Results are expressed as normalized log (mean gene activity). Hierarchical clustering is done according to the NT dataset; 5 clusters are identified: Cl_1, Cl_2, Cl_3, Cl_4, Cl_5.

### Figure S3 (related to Figure 3)

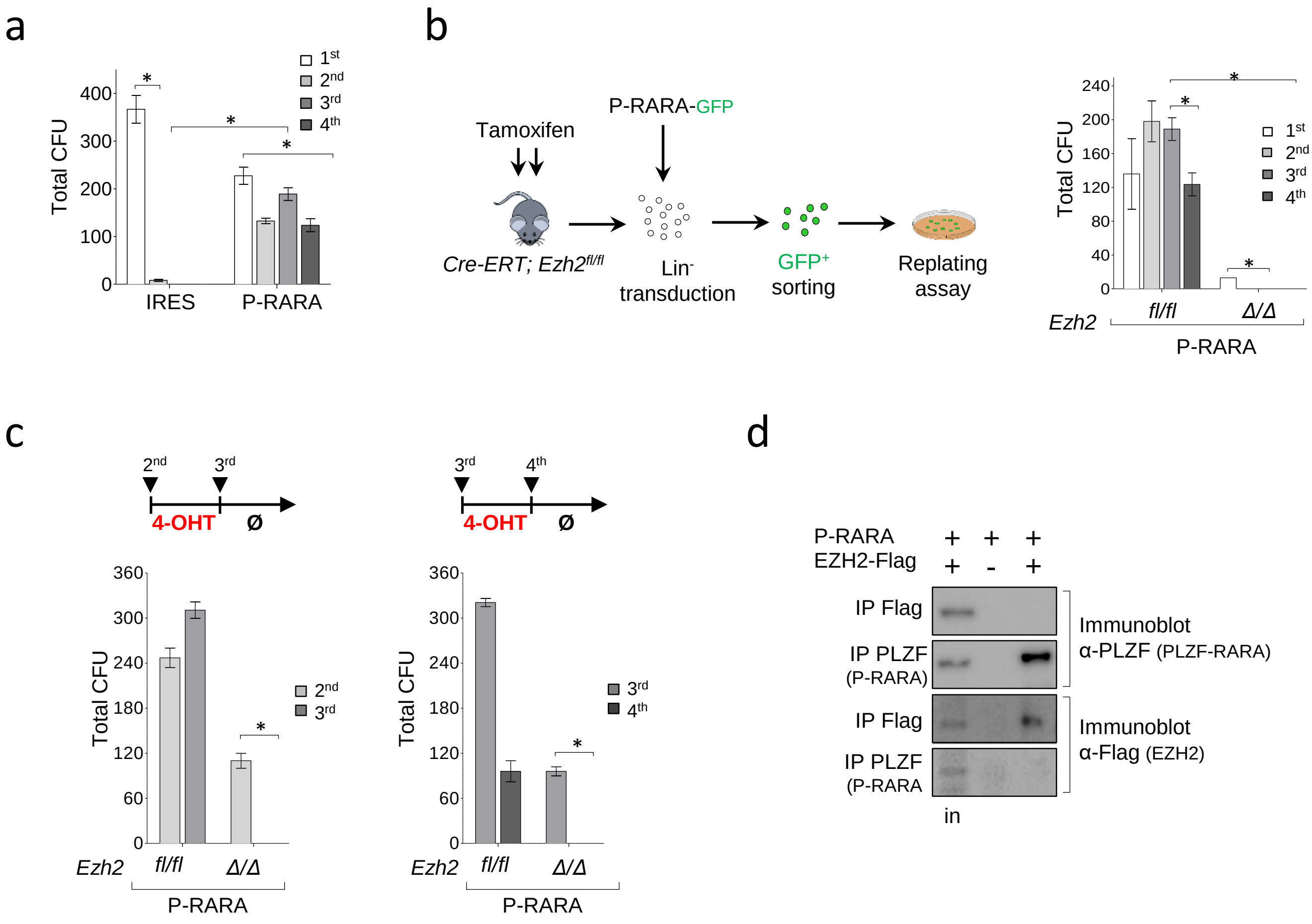


**Figure S3 related to Figure 3. EZH2 relevance in PLZF-RARA transformation. (A)** PLZF-RARA transformation efficiency evaluated by replating assay. Total Colony Formation Units (CFU) are counted at each replating step. IRES: cells transduced with an empty vector; P-RARA: cells transduced with PLZF-RARA. Results are expressed as a mean SD of three experiments (n=3). *p-value < 0.05. (**B**) Impact of Ezh2 loss in PLZF-RARA transformation. On the left, experimental scheme of Ezh2 deletion *in vivo*. *Cre-ERT;Ezh2$^{fl/fl}$* mice are intraperitoneally injected with 100 µl of tamoxifen dissolved in corn oil at a concentration of 10 mg/ml for 5 consecutive days. Lineage negative (Lin$^-$) cells are purified and transduced with an *empty-IRES-GFP* (IRES) or a *PLZF-RARA-IRES-GFP* (P-RARA) retroviral construct. Lin$^-$ GFP positive cells are FACS sorted, seeded into methylcellulose and replated. On the right, total Colony Formation Units (CFU) are counted at each replating step. IRES: cells transduced with an empty vector; P-RARA: cells transduced with PLZF-RARA. Results are expressed as a mean SD of three experiments (n=3). *p-value < 0.05. (**C**) Impact of EZH2 loss in PLZF-RARA maintenance. *Ezh2* deletion in Lin$^-$ GFP positive cells is obtained by adding 150nM 4-OHT in the methylcellulose at the 2$^{nd}$ (left panel) or the 3$^{rd}$ (right panel) round of plating. Replating efficiency is monitored by counting the total Colony Forming Units (CFU) of transformed (P-RARA). Results are expressed as a mean SD of three experiments (n=3). *p-value < 0.05. (**D**) Nuclear extracts of 293T cells co-transfected with PLZF-RARA (P-RARA) and EZH2-Flag immunoprecipated with an anti-Flag or an anti-PLZF. IPs are immunoblotted with an anti-PLZF (upper panel) or an anti-Flag antibody (lower panel). Inputs (in) represent 2% of samples processed in each IP.

### Figure S4 (related to Figure 4)

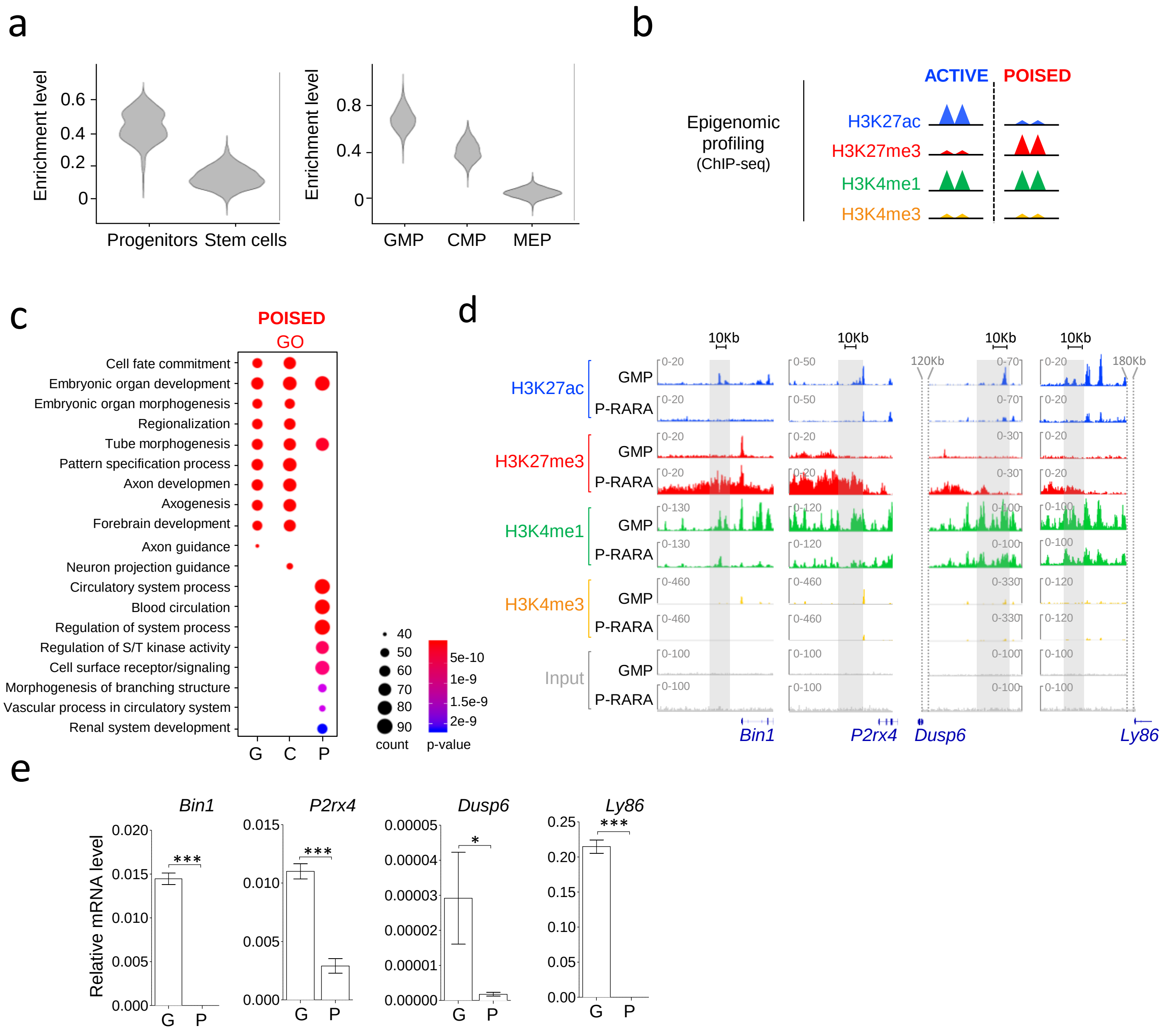


**Figure S4 related to Figure 4.** (**A**) Left panel: Violin plots showing hematopoietic progenitor (C-Kit$^+$, Sca1$^-$) and stem cell (C-Kit$^+$, Sca1$^+$) signatures in the NT scRNA-seq dataset. Right panel: Violin plots showing Granulocyte-Monocyte Progenitor (GMP: C-Kit$^+$, Sca1$^-$, Cd34$^+$, FcγR$^+$), Common Myeloid Progenitor (CMP: C-Kit$^+$, Sca1$^-$, Cd34$^{low}$, FcγR$^{low}$) and Megakaryocyte-Erythrocyte Progenitor (MEP: C-Kit$^+$, Sca1$^-$, Cd34$^-$, FcγR$^-$) in the NT scRNA-seq dataset. The signature score represents the global expression of annotated genes for each signature extracted from Nestorowa et al. (Nestorowa et *al*., 2016). (**B**) Schematic representation of active and poised enhancers. Epigenetic marks (H3K27ac, H3K27me3, H3K4me1, H3K4me3) are monitored by ChIP-sequencing (ChIP-seq). Enhancer regions are determined with H3K4me1 (high) and H3K4me3 (low) levels. Active enhancers are H3K4me1+, H3K4me3-, H3K27ac+, H3K27me3-. Poised enhancers are H3K4me1+, H3K4me3-, H3K27ac- H3K27me3+. Triangles represent the histone marks of interest. (**C**) Gene Ontology (GO) associated with poised enhancers using ClusterProfiler package tools. Circle size represents the number of genes observed in the GO term associated with poised enhancers. Color scale values indicate the p-value. G (Enhancers specific for GMP condition), C (Enhancers shared by both conditions), P (Enhancers specific for PLZF-RARA condition). (**D**) Representative Integrative Genomics Viewer (IGV) tracks of H3K27ac, H3K27me3, H3K4me1 and H3K4me3 in P-RARA-transformed progenitors and GMPs at enhancer regions that switch from an active (in GMP) to poised (in P-RARA) state. The gray box underlines the region of interest. (**E**) mRNA levels of switched enhancer nearby genes in GMP (G) and PLZF-RARA (P) conditions monitored by qPCR. mRNA values are normalized to *β2microglobulin* and are expressed as a mean SD of three independent experiments (n=6). *p-value < 0.05, *** p-value < 0.001.

### Figure S5 (related to Figure 4)

a

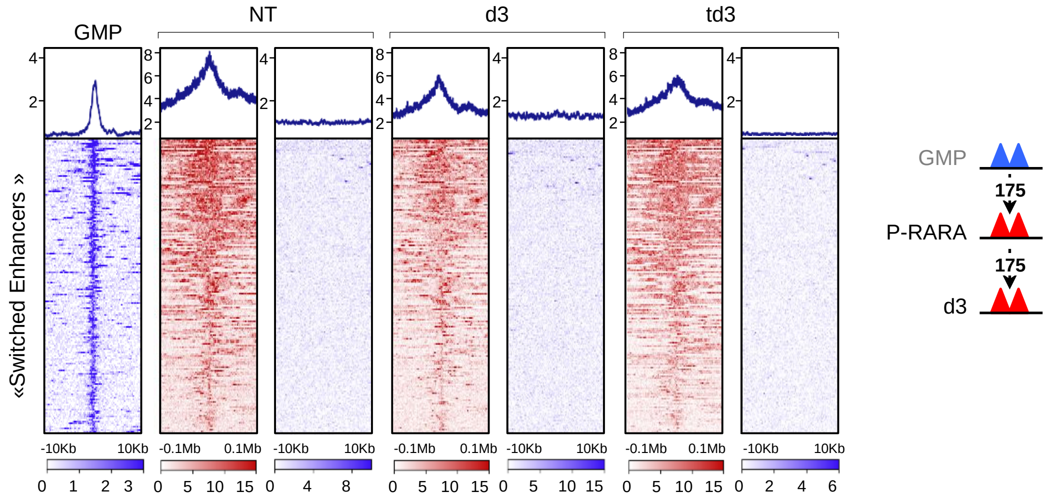


b

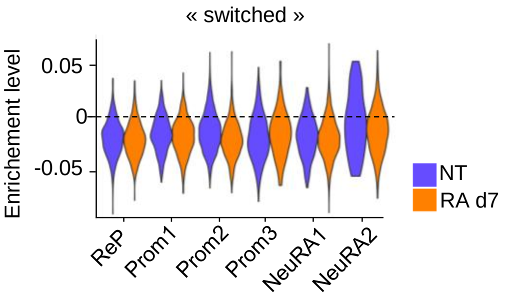


**Figure S5 related to Figure 4.** (**A**) Left panel: Plot Heatmap of H3K27ac and H3K27me3 signals in GMP, Crtl, d3 and td3 conditions at enhancers that switch from active (in GMP) to poised state (in PLZF-RARA). H3K27ac and H3K27me3 signals are plotted 10Kb and 0.1Mb upstream and downstream the enhancer center. Right panel: Active enhancers are represented by blue triangles and poised enhancers by red triangles. (**B**) Violin plot showing “Enhancer switch” signature scores per treatment in each cluster identified by scRNA-seq in Figure 1. Signature score represents the global expression of annotated genes for the selected signature.

### Figure S6 (related to Figure 5)

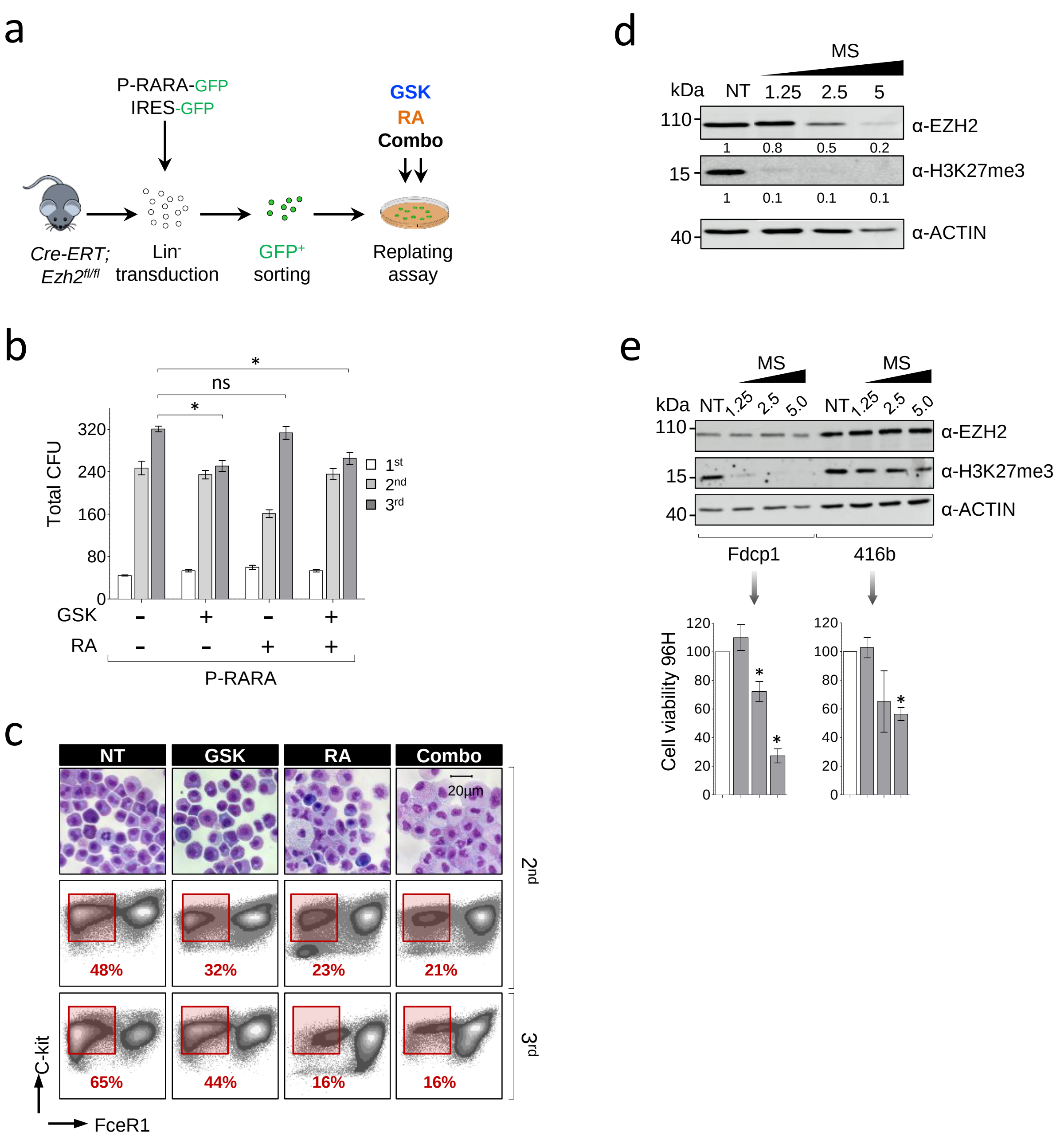


**Figure S6 related to Figure 5.** (**A**) For Ezh2 catalytic inhibition and to monitor the effect of RA treatment on this *ex-vivo* model, Lin- GFP positive cells are seeded in methylcellulose containing 2.5µM GSK126 (GSK) or 1µM Retinoic Acid (RA) or both drugs (Combo). Drugs are added at each round of plating. Cells are re-plated every 1-2 weeks (**B**) Replating efficiency of transduced PLZF-RARA cells (P/RARA) with different treatments is monitored by counting the total Colony Forming Units (CFU). Results are expressed as a mean SD of three experiments (n=3). ns: not significant, *p-value < 0.05. (**C**) Cell differentiation at the $2^{nd}$ and $3^{rd}$ rounds of plating evaluated by May Grunwald Giemsa (MGG) staining (magnification 64X; bar 20 µm) (upper panel) and FACS analysis (middle and lower panels; C-Kit and FceR1 markers are monitored). (**D**) *In vitro* efficiency of MS1943 on EZH2 protein level. PLZF-RARA TG cells are treated for 96H with increasing doses of MS1943 (MS) (NT: 0, 1.25, 2.5, 5 µM). Global levels of Ezh2 and H3K27me3 detected by western blotting (WB) in each indicated condition. Actin is used as loading control. Signal intensity is measured and normalized according to the loading control and to the untreated condition. **(E)** Impact of MS1943 on murine myeloid cell lines. Fdcp1 and 416b are treated for 96H with increasing doses of MS1943 (MS) (NT: 0, 1.25, 2.5, 5 µM). Upper panel: Global levels of Ezh2 and H3K27me3 detected by western blotting (WB) in each indicated condition. Actin is used as loading control. Lower panel: Cell viability monitored by bioluminescence, *p-value < 0.05.
